# A synthetic peptide rescues rat cortical neurons from anesthetic-induced cell death and modulation of growth and synaptic assembly

**DOI:** 10.1101/2020.06.18.159889

**Authors:** Fahad Iqbal, Marcus Pehar, Andrew J. Thompson, Urva Azeem, Kiana Jahanbakhsh, Nerea Jimenez-Tellez, Rasha Sabouny, Shadab Batool, Atika Syeda, Jennifer Chow, Pranav Machiraju, Timothy Shutt, Kamran Yusuf, Jane Shearer, Tiffany Rice, Naweed I. Syed

**Affiliations:** University of Calgary; University of Oxford

**Author notes:** indicates equal contribution (Marcus Pehar, Andrew J. Thompson, Urva Azeem, Kiana Jahanbakhsh). indicates corresponding author **Corresponding Author:** Naweed I. Syed **Email Address of Corresponding Author:**. **Postal Code:** T2N 4N1.

**Keywords:** Anesthetics, Neurotoxicity, P110, Desflurane, Propofol, Ketamine, Neuroprotection, Mitochondria, Synapse

## Abstract

Anesthetics, although necessary for many modern procedures, exert neurotoxic effects in various experimental models; the underlying mechanisms, however, remain unknown. In the absence of this information, any *in vivo* mitigation strategy would be challenging. To help address these gaps, we sought to determine whether preserving mitochondrial network integrity with a non-toxic, short-life synthetic peptide, P110, could protect cortical neurons against both inhalational and intravenous anesthetic-induced neurotoxicity. This study provides the first direct comparative account of three key general anesthetics (desflurane, propofol, and ketamine) under identical conditions and demonstrates their impact on cellular viability, neurite outgrowth, and synaptic network assembly on neonatal rat cortical neurons. Further, we discovered that the inhibition of Fis1 receptor-mediated fission reverses anesthetic-induced aberrations in an agent-specific manner. Our data thus underscore the importance of a reductionist approach, offering unique opportunities to explore non-toxic, mitochondria-based therapeutic tools for mitigating anesthetic-induced harm to the developing brain.

## Introduction

Anesthetics are required for most major clinical procedures as they fulfill numerous surgical needs. Recent studies have however, shown that a majority of clinically used anesthetics also exert significant long-term, neurotoxic effects in both animal models and human neural stem cells. The evidence of harm ranges from cell death to suppression of neurite outgrowth to perturbation of synaptic connectivity^1–3^. Moreover, some epidemiological studies have also implicated a variety of anesthetic agents in learning, memory and cognitive dysfunction in children, although the clinical evidence remains equivocal^4,5^. As a result, concerns about the long-term effects of anesthetics have arisen – especially when administered to young children or pregnant mothers^6,7^.

Although our understanding of the mechanisms of how various anesthetics function has vastly improved in recent years^4,8,9^, a complete picture of their modes of action in the context of the developing brain remains unclear - owing mainly to the complexity of the human brain and the ethical implications of using humans as experimental subjects. Moreover, the behavioral learning and memory deficits observed in freely moving animals exposed to anesthetics is similarly irreconcilable at the cellular and molecular levels. As such, clinicians are left with conflicting data from both human subjects and animal models. Notwithstanding the abundance of anecdotal evidence from human studies suggesting that anesthetic exposure may affect learning, memory and cognitive function in both young children and the elderly alike, the evidence remains however, largely anecdotal^4,10^. In the absence of definitive answers regarding the potential sites of anesthetic action and the nature of their harmful effects, strategies to mitigate their long-term side effects cannot be put in place. Moreover, a direct comparison between γ-aminobutyric acid (GABA)-ergic and N-Methyl-D-aspartic acid (NMDA)-ergic general anesthetics (the major targets for most anesthetics), and the extent to which they compromise neuronal development is yet to be performed.

To provide a comprehensive and comparative account of various anesthetics’ actions, we examined one inhalational anesthetic, desflurane, and two intravenous agents, propofol, and ketamine. Desflurane acts primarily by promoting inhibitory GABAergic synaptic transmission^4^, whereas propofol is the most commonly used clinical intravenous agent, and like desflurane, it acts primarily via GABA agonism^4^. Ketamine on the other hand, is an intravenous psychedelic compound as well as an anesthetic; it acts primarily by blocking NMDA receptor-mediated excitatory synaptic transmission^4^. We examined the above agents under identical and clinically relevant conditions for their differential effects on cell viability, neurite outgrowth, and synapse formation in cortical rat neurons and identified potential sites for their harmful effects. Recent studies have implicated mitochondrial dysfunction, specifically fragmentation of its networks as a common factor in several degenerative diseases^11–14^, and most recently, mitochondrial dysfunction vis-à-vis anesthetic-induced aberrations has also come to light^2,15–17^. We therefore sought to identify how these agents affect mitochondria and its function and determined whether these detrimental effects could be prevented by using a synthetic peptide, P110, which restores mitochondrial networks in stressed neurons.

## Methods

### Ethical Approval

All animal procedures were carried out in accordance with the standards established by the University of Calgary Animal Care and Use Policy under the Canadian Council on Animal Care. All experimental protocols have been approved by the University of Calgary Animal Care Committee. All findings were obtained without unnecessary pain and suffering of animals and follow current best practice and use most appropriate species for the study^18^. The protocols used are largely identical to those used in our previous study^2^. These have been adapted and reproduced here to reflect the experiments carried out in this study.

Both male and female postnatal day 0 wild-type Sprague Dawley strain code 400 rats from Charles River Laboratories were used. The mothers of these postnatal day 0 rats were kept in a conventional room set at 21°C on a 12-hour light/dark cycle from 7:00am to 7:00pm local time and were fed *ad libitum*. Sedation was achieved via ice induced hypothermia, where the neonatal rat was wrapped in tissue paper and placed in a chamber filled with ice for 10 minutes^19^. Decapitation was performed following loss of movement, prior to the regaining of consciousness. Following decapitation, cortical tissue was collected.

### Primary rat neuronal cell culture and P110 treatment

Sprague-Dawley rat frontal cortexes were isolated from postnatal day 0 pups and enzymatically treated with papain (50 u/mL; Worthington Biochemical Corp., Lakewood, NJ, USA). The cells were then triturated in glass pipettes of decreasing size to create a uniform cell suspension and plated at appropriate confluence onto cover slips coated with laminin (2 μg/mL; MilliporeSigma, St. Louis, MO, USA) and poly-D-lysine (30 μg/mL; MilliporeSigma, St. Louis, MO, USA) in culture dishes. Cells settled for one hour at 37°C and 5% CO_2_ prior to the addition of additional 2mL culture media. The culture medium used was Neurobasal medium supplemented with 2% B27 (Thermo Fisher Scientific, Waltham, MA, USA), L-Glutamine (200 mM; Thermo Fisher Scientific, Waltham, MA, USA), 4% FBS (Thermo Fisher Scientific, Waltham, MA, USA), and penicillin-streptomycin (10 000u/mL penicillin, 10 000μ/mL streptomycin; Invitrogen, Waltham, MA, USA), which was changed (50 % removed and replaced) every 3 days. Some culture dishes were treated with 1uM P110 (provided by Daria Mochly-Rosen, Department of Chemical and Systems Biology, Stanford University School of Medicine, Stanford, CA, USA) for one hour in culture media, while controls had normal culture media without any P110.

The cells were incubated at 37°C and 5% CO_2_ for 1 hour before anesthetic exposure/treatment.

### Desflurane exposure

After 1-hour incubation at 37°C and 5% CO_2_, the cells were exposed to 0.5 minimum alveolar concentration (MAC) equivalent of desflurane (4.3 Vol%; Baxter Corporation, Mississauga, ON, Canada) in an airtight modular incubator chamber (Billups-Rothenberg) for one hour. Desflurane-medical air gas mixtures were vaporized using a Datex-Ohmeda Aestiva/5 vaporizer and concentrations were monitored with a GE Healthcare Gas Analyzer. Controls were exposed to medical air only. After 1 hour of desflurane-medical air mixture or just medical air exposure, the neurons were placed back and maintained in an incubator (37°C, 5% CO_2_) until use.

### Propofol and ketamine treatment

After 1-hour incubation at 37°C and 5% CO_2_, propofol and ketamine were added to the experimental treatments individually at appropriate concentrations (10μM propofol and 5μM ketamine, respectively). The choice for these concentrations was based on previous studies wherein they were the lowest end to have reliably affected neuronal characteristics^15,20,21^. These drugs were then diluted out via half media replacement the next day. The cells were maintained in an incubator (37°C, 5% CO_2_) until use.

### Live-cell fluorescent imaging and confocal microscopy

To assess the impact of desflurane on the morphological integrity of the mitochondria, cells were grown for 4 days post-culture and then had their media replaced with media containing MitoTracker™ Red CMXRos (ThermoFisher Scientific, cat. M7512) (200 nM) and incubated with the cells for 20 mins at 37°C and 5% CO_2_. The cells were then washed 3 times with warmed DPBS, and fresh media was added. The cells were then imaged.

To simultaneously assess the impact of anesthetics on mitochondrial morphology and SO production, cells were grown for 4 days post-culture and then had their media replaced with media containing MitoTracker™ Green (ThermoFisher Scientific, cat. M7514) (70nM) and MitoSOX™ Red (ThermoFisher Scientific, cat. M36008) (5μM) and incubated with the cells for 20 mins at 37°C and 5% CO_2_. The cells were then washed 3 times with warmed DPBS, and fresh media was added. The cells were then imaged.

Cells were imaged at this time point to minimize glial interference that would complicate neuron identification. Mitochondrial morphology was quantified by assessing the percentage of cells that exhibited a particular morphology throughout three biological replicates. Each cell was classified as having a given mitochondrial morphology based on the predominant mitochondrial morphology present in each cell according to the following criteria ^2^: mitochondria were considered fragmented when less than 0.75 μm in length, intermediate when 0.75 μm to 3 μm in length, and fused when greater than 3 μm in length. These characterizations were reflected on a three-point fragmentation scale and cells were classified as having fused, intermediate, or fragmented mitochondrial networks **(Figure 1)**.

**Figure 1:**
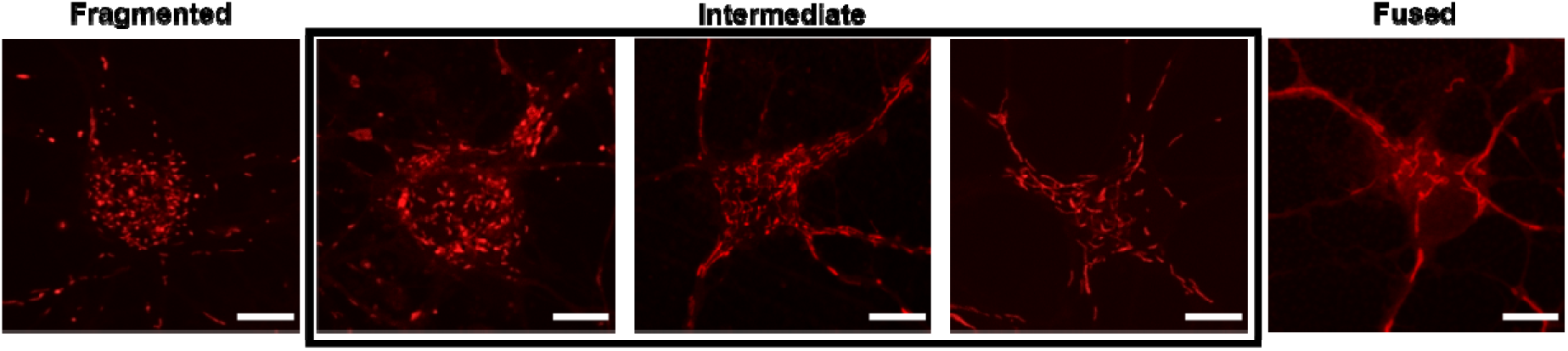
Mitochondrial morphology quantification reference scale. Cells were manually classified into one of 3 categories. Cells which exhibited more extreme fragmentation or hyper-fusion were counted to their closest equivalent. Neurons labeled with Mitotracker™ Red CMXRos. Scale bars indicate 10μm.

Fluorescence images were taken with an Olympus SD-OSR spinning-disk confocal microscope with a mounted incubation system (Olympus Corp.) using the same imaging parameters for every dish. A 100x objective was used. Mitochondrial SO production was measured by calculating the mean grey intensity of SO signal per field of view and compared between treatments using the same imaging parameters and thresholds in ImageJ. The intensity of SO signal that localized to the MitoTracker™ Green signal was also calculated and compared in the same manner.

### Cell viability assay

The effects of anesthetics on neuronal viability were tested 3 days post-culture via the LIVE/DEAD® Viability⁄Cytotoxicity Kit (Molecular Probes). This time point was selected to minimize any glial proliferation that could mask any changes in viability of developing neurons. Specifically, the cells were exposed to calcein-AM (green, live cells) and ethidium homodimer-1 (red, dead cells) dyes at room temperature for 15 min and imaged using a Zeiss Axio Observer Z1 microscope (Zeiss Corp.) with a 10x objective. The percentage of alive cells were automatically counted using the Cell Counter plugin in ImageJ with the same thresholds for all the treatments.

### Immunofluorescent staining

Immunofluorescent staining to assess the impact of anesthetics on synaptic network assembly was conducted on neurons grown for 10 days post-culture. Fixation was performed with 4% paraformaldehyde for 20 mins at room temperature. The fixed cells were permeabilized for 1 hour at room temperature with incubation media consisting of 0.1% Triton X with 5% goat serum. A primary rabbit monoclonal antibody for synaptophysin (1:500) (Abcam; EP1098Y) and mouse monoclonal antibody for PSD-95 (1:1000) (NeuroMab; 75-028) were applied to the cultures overnight at 4°C. Secondary antibodies of AlexaFluor 561 goat anti-rabbit IgG (1:400) (Invitrogen; A11011), and AlexaFluor 488 goat anti-mouse IgG (1:400) (Invitrogen; A28175) were then applied. Dishes were mounted with MOWIOL mounting media with DAPI to stain nuclei (Sigma-Aldrich). Cell were imaged with a Zeiss Axio Observer Z1 microscope (Zeiss Corp.) with a 63x objective. Synaptic puncta density was quantified manually with a focus on primary neurites which emanated either from the cell body or secondary projections from other dendritic neurites under identical thresholds. Multiple processes from a minimum of 5 cells per biological replicate were quantified.

To assess the impact of anesthetics on neuronal cytoskeletal growth, the neurons were stained with primary chicken polyclonal antibody against the 160 kDa fragment of neurofilament (1:500) (Novus Biologicals; NB300-222) and the secondary antibody of AlexaFluor 488 goat anti-chicken IgY (1:400) (Invitrogen; A32931). The cells were imaged using a Zeiss Axio Observer Z1 microscope (Zeiss Corp.) with a 20x objective. Image acquisition parameters (laser intensity, pinhole sizes, exposure times, gain settings etc.) were kept consistent throughout all treatments. Average neurite outgrowth was quantified using the ImageJ plugin NeuronJ (minimum n◻=◻19 areas of 0.4 mm^2^ for each treatment) with identical conditions and thresholds for each condition.

Antibody specificity data has been previously published via negative control experiments that show no immunofluorescence in the absence of primary antibodies^2,22^.

### Statistical analysis

Statistical significance tests were conducted with GraphPad Prism 8. Two-tailed t-tests were conducted to compare between two independent means. One-way ANOVA was used to compare groups with one independent variable (ie. control vs. propofol vs. ketamine). Two-way ANOVA was used to compare groups with two independent variables (ie. P110 treatment and anesthetic treatment or exposure). Tukey’s or Sidak’s multiple comparisons tests were conducted as post-hoc tests as appropriate. Values were graphed as mean◻±◻standard deviation (SD) across at least 3 biological replicates. To independently validate the quantification and increase confidence in the observed trends across separate experiments, at least 2 blind counts were conducted per set of experiments. Differences between data were considered significant if appropriate post-hoc statistical tests resulted in *p* ≤ 0.05. Significance parameters were indicated with asterisks as follows: \**p* ≤ 0.05, \*\**p* ≤ 0.01, \*\*\**p* ≤ 0.001, \*\*\*\**p* ≤ 0.0001.

#### Data Availability

The data that support the findings of this study are openly available in the Mendeley Data Depository at doi:10.17632/5j76t2b374.1.

## Results

### Desflurane, propofol, and ketamine cause significant cell death in primary rat cortical neurons

To directly assess the effects of both GABAergic and NMDAergic anesthetics on neuronal health, and to resolve varied interpretations emanating from conflicting results in the literature, we tested the effects of desflurane (4.3 Vol.%), propofol (10μM), and ketamine (5μM) on cortical neuronal viability under identical experimental conditions. This clinically relevant concentration for desflurane equates to 0.5 minimum alveolar concentration (MAC). The concentrations for the comparative agents propofol and ketamine were chosen based on their clinical utility and as per the previously published studies in various animal models^15,20,21^.

Cells cultured under control conditions displayed healthy, branching neurites with there being no signs of vacuolation **(Figure 1; 1a and 2a)**. On the other hand, neurons exposed to desflurane **(Figure 2; 1b)**, propofol **(Figure 2; 2b),**and ketamine **(Figure 2; 2c)**exhibited signs of vacuolation. A quantitative analysis of these observations was conducted via live/dead assay and multiple areas (0.58 mm^2^) of three replicates of each condition were imaged and quantified under identical thresholds. Examples of dead cells are indicated with asterisks. Two-tailed t-tests confirmed that desflurane-exposed cells exhibited significantly reduced cell viability: (*Mean* = 72.38%, *SD* = 6.99% alive, *p* = 0.0346, n = 53 areas) as compared to control (*M* = 88.13%, *SD* = 3.62% alive, n = 35 areas) **(Figure 2; 1e)**. Similar observations were made for the intravenous agents, which showed that propofol-treated cells exhibited increased cell death (*M* = 74.09%, *SD* = 1.67% alive, *p* < 0.0001, n = 42 areas) as compared to control (*M* = 89.15%, *SD* = 3.89% alive, n = 54 areas) **(Figure 2; 2g)**. Ketamine also caused cell death compared to control (M = 80.88%, SD = 1.88% alive, *p* = 0.0032, n = 12 areas), although not to the level of propofol (*p* = 0.0169) (**Figure 2; 2g)**. Taken together, these data demonstrate that the anesthetics desflurane, propofol, and ketamine, all cause significant apoptosis in rat cortical rat neurons.

**Figure 2:**
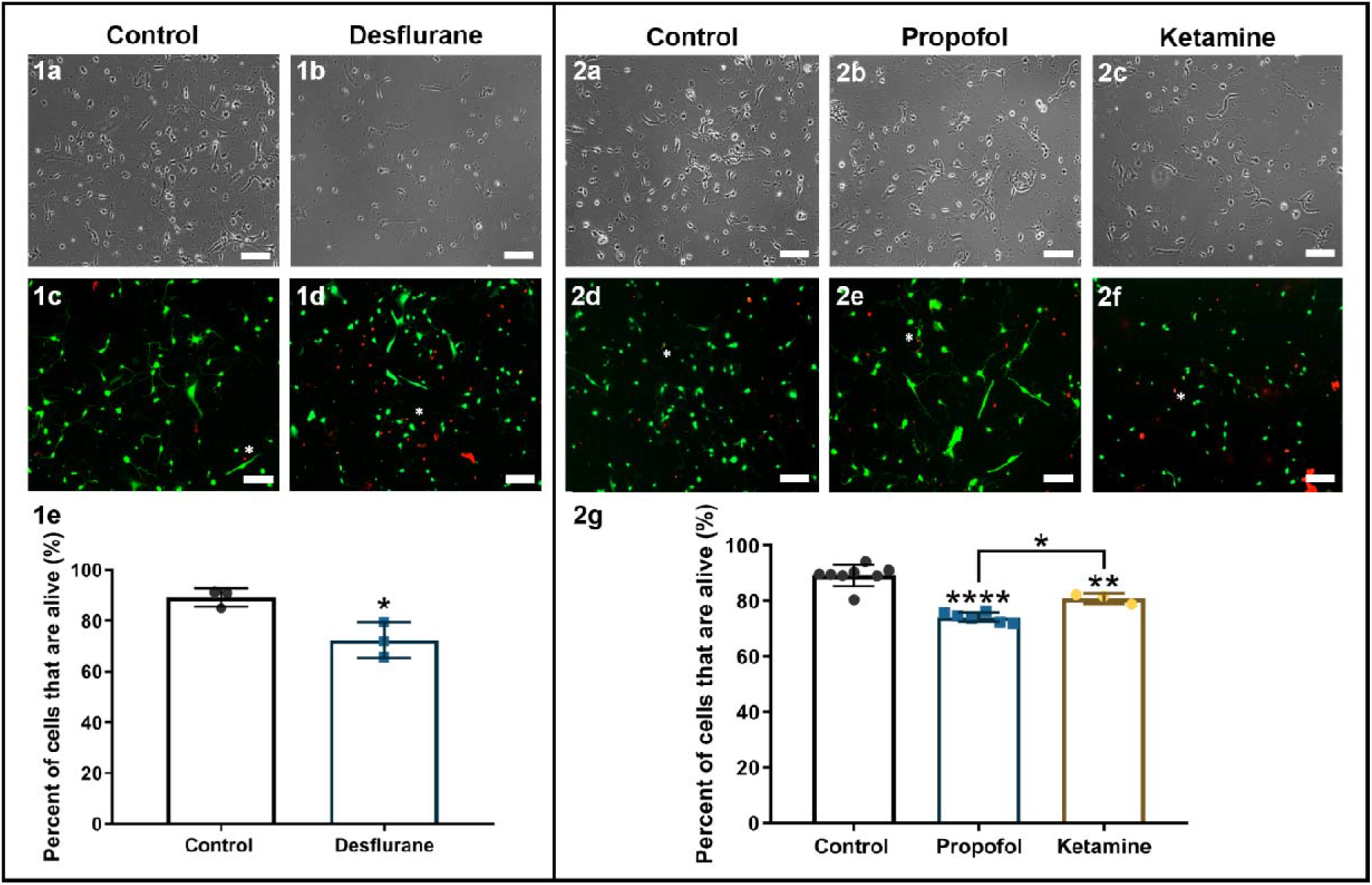
Anesthetics-induced apoptosis of primary cortical neurons. Representative phase-contrast images depicting different experimental conditions **(1a,1b,2a,2b,2c)**. Representative live-fluorescent images of different conditions labeled with a LIVE/DEAD™ Viability/Cytotoxicity Assay where live cells are identified with calcein-AM (green) and dead cells are identified with ethidium homodimer-1 (red) **(1c,1d,2d,2e,2f)**. Quantification of percentage of live cells **(1e and 2g)**. Exposure to 4.3Vol% desflurane, treatment with 10μM propofol, or treatment with 5μM ketamine caused significant cell death, although ketamine was not as detrimental as propofol. Two-tailed t-test. * *p* < 0.05, ** *p* < 0.01, **** *p* < 0.0001. Bars indicate ±SD measured across biological replicates. Each data-point indicates the mean viability reported in each biological replicate. Phase and fluorescent images are for different regions. Scale bars indicate 100μm.

### Anesthetics differentially affect neurite outgrowth and synaptic network assembly in developing rat cortical neurons

#### Neurite outgrowth

Developing neurons extend their axonal and dendritic processes and branch out to reach their synaptic targets; failure to do so results in perturbed connectivity and compromised functional assembly of the brain. To assess whether general anesthetics with different modes of action affect neurite growth of surviving neurons, we next tested the effects of desflurane, propofol, and ketamine on the ability of developing neurons to exhibit outgrowth. For this, the cultured neurons were stained with polyclonal antibodies against the 160 kDa fragment of neurofilament **(Figure 3A)** and average neurite lengths were quantified in multiple areas of 0.15 mm^2^. We found that neurons exposed to 4.3 Vol% desflurane exhibited neurite lengths that were not significantly lower than control (*M* = 0.70, *SD* = 0.09 fraction of control, *p* = 0.0599, n = 53 areas) compared to control (*M* = 1.00, *SD* = 0.16, n = 40 areas) (**Figure 3C)**. Neurons treated with 10μM propofol also showed no significant change in neurite outgrowth (*M* = 0.82, *SD* = 0.05 fraction of control, *p* = 0.4386, n = 31 areas) compared to control (*M* = 1.00, *SD* = 0.13, n = 62 areas) **(Figure 3C)**. Neurons treated with ketamine however, exhibited more robust growth than their control counterparts (*M* = 1.91, *SD* = 0.36 fraction of control, *p* = 0.0003, n = 19 areas) and compared to those treated with propofol alone (*p* = 0.0002) **(Figure 3C)**. Taken together, these data demonstrate that neither desflurane nor propofol inhibit average neurite outgrowth length per cell, whereas ketamine enhances neuronal outgrowth of cortical neurons.

**Figure 3:**
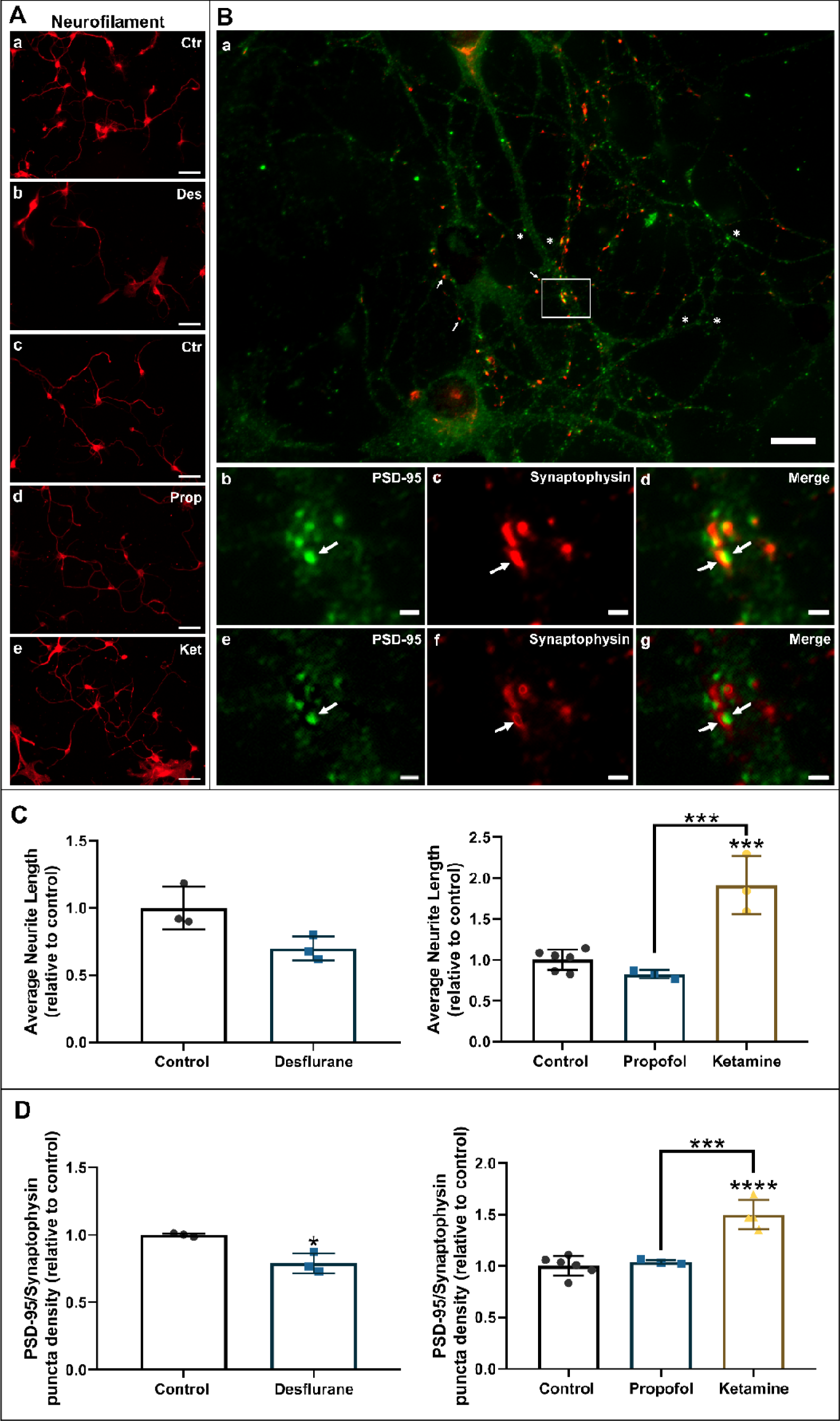
Desflurane but not propofol decreases synaptic protein expression whereas ketamine increased both neurite outgrowth and synaptic protein expression. Representative fluorescent images of cortical neurons stained for neurofilament **(A)**. Scale bars indicate 50μm. Representative fluorescent image that is labelled for the post-synaptic marker PSD-95 (green) and the presynaptic marker synaptophysin (red) **(B)**. Close-up of a synaptic punctum where the juxtaposition of PSD-95 **(b)** and synaptophysin **(c)** produces a characteristic yellow, which was quantified as a punctum **(d)**. A sharpened image is shown in the final row to verify and display the localization of these signals at the synaptic site **(e-g)**. Average puncta density per neurite was manually quantified as number of puncta per unit length wherever these characters for a synapse were met. Scale bars indicate 10μm **(a-d)** and 1μm **(b-g)**. Quantification of average neurite length relative to control **(C)**. Exposure to 4.3 Vol.% desflurane did not decrease average neurite lengths. Treatment with 10μM propofol did not significantly affect neurite outgrowth. Treatment with 5μM ketamine resulted in excessive growth compared to control. Quantification of average PSD-95/synaptophysin puncta per unit neurite length relative to control **(D)**. Exposure to 4.3 Vol.% desflurane decreased average synaptic puncta density. Treatment with 10μM propofol did not affect average synaptic puncta density. Treatment with 5μM ketamine increased average puncta density. Two-tailed t-test or one-way ANOVA. * *p* < 0.05, ** *p* < 0.01, *** *p*◻<◻0.001 as determined by pairwise comparison of post-hoc tests. Bars indicate ±SD measured across biological replicates. Each data-point indicates the mean average neurite lengths, and mean puncta density per biological replicate.

#### Synaptic network assembly

Robust synaptic network assembly is prerequisite to proper brain development. To examine the effects of various anesthetics on synaptic network assembly, cells were stained for two synaptic proteins – synaptophysin, a presynaptic vesicle protein, and postsynaptic density protein (PSD-95); their expression patterns and colocalized juxtapositions were taken as an index of synaptic specialization **(Figure 3B)**. Only puncta that met these requirements were counted, such as the example highlighted in **Figures 3B (b-g)**. Instances where there was only synaptophysin signal (examples indicated with an arrow in **Figure 3Ba**), or only PSD-95 signal (examples indicated with an asterisk in **Figure 3Ba**) with no juxtaposition to each other and no overlapping yellow signal were not considered and indicators of a synapse. Neurons cultured under control conditions exhibited uniform expression of PSD-95 (green) with punctate juxtaposed against synaptophysin (red). Desflurane-exposed cells exhibited decreased density of synaptic puncta (*M* = 0.79, *SD* =◻0.07 fraction of control, *p*◻= 0.0356, n◻=◻53 neurites) compared to control (*M* = 1.00, *SD* =◻0.01 fraction of control, n◻=◻131 neurites) **(Figure 3D)**. Propofol, however, did not affect synaptic puncta density (*M* = 1.04, *SD* = 0.02 fraction of control, *p* = 0.8635, n = 65 neurites) compared to control (*M* = 1.00, *SD* = 0.1 fraction of control, n = 112 neurites) **(Figure 3D)**. Interestingly, ketamine-treated cells exhibited increased synaptic puncta density compared to control (*M* = 1.5, *SD* =◻0.14 fraction of control, *p*◻< 0.0001, n◻=◻81 neurites) and compared to those treated with propofol alone (*p* = 0.0004) **(Figure 3D)**. These data thus show that desflurane decreases synaptic puncta formation, whereas propofol does not significantly affect synaptic assembly. Contrary to the GABAergic desflurane, ketamine increased synaptic puncta density as revealed by the expression of two key synaptic markers, synaptophysin and PSD-95, in cortical neurons.

### Propofol and ketamine compromise mitochondrial networks whereas ketamine elevates mitochondrial superoxide production

All cellular processes depend on underlying energy dynamics, and thus the mitochondria have become the centre of extensive investigation as a potential target site for anesthetic-induced neurotoxicity and neurodegeneration. To assess the relative effects of the two intravenous agents propofol and ketamine on mitochondrial morphology, and to determine if any of the observed changes were accompanied by functional deficits, we examined mitochondrial superoxide (SO) production, which is an indicator of mitochondrial and cellular health. We labeled the neurons with MitoTracker™ Green FM for mitochondria and MitoSOX™ Red for SO production and discovered that under control conditions, mitochondria exhibited typical, elongated, rod-like morphology **(Figure 4A).** Conversely, neurons exposed to anesthetics exhibited small, round morphology, with fewer elongated rod-like mitochondria, which is a characteristic for fragmented mitochondrial networks and mitochondrial toxicity, dysfunction, and the invocation of intrinsic apoptotic pathway **(Figure 4A)**. To validate these observations, we quantified the mitochondrial morphology in cortical neurons using a 3-point scale (fragmented, intermediate, and fused) **(Figure 1)** and assessed the data via two-way ANOVA. We found that both propofol (*M* = 50.99%, *SD* = 7.65%, *p* < 0.0001, n = 115 cells) and ketamine (*M* = 44.97%, *SD* = 5.68%, *p* = 0.0008, n = 84 cells) caused aberrant mitochondrial fragmentation compared to control (*M* = 24.87%, *SD* = 3.45%, n = 136 cells), with these conditions showcasing a higher proportion of cells with fragmented mitochondrial networks **(Figure 4B)**. We concomitantly examined SO production under these experimental conditions to quantify potential differences in the effects propofol and ketamine on mitochondrial function. By measuring fluorescence intensity of MitoSOX™ Red over the full field of view, we found that ketamine (*M* = 213.19%, *SD* = 55.35% of control, *p* = 0.0059 compared to control, n = 49 areas), but not propofol (*M* = 152.1%, *SD* = 17.68% of control, *p* = 0.1736 compared to control, *p* = 0.146 compared to ketamine alone, n = 52 areas) and caused a dramatic increase in SO production in the cells compared to control (*M* = 100%, *SD* = 29.13% of control, n = 85 areas) **(Figure 4C)**. We further investigated this relationship and quantified superoxide signal that colocalized with the mitochondrial signal. We found similar trends for both propofol (*M* = 156.3%, *SD* = 46.17% of control, *p* = 0.5981, n = 52 areas) and ketamine (*M* = 275%, *SD* = 114.7% of control, *p* = 0.0348 compared to control and *p* = 0.2051 compared to propofol alone, n = 49 areas) compared to control (*M* = 100%, *SD* = 65.42% of control, n = 85 areas) **(Figure 4D)**. These data thus show both propofol and ketamine cause a robust increase in mitochondrial fragmentation and ketamine increases SO production.

**Figure 4:**
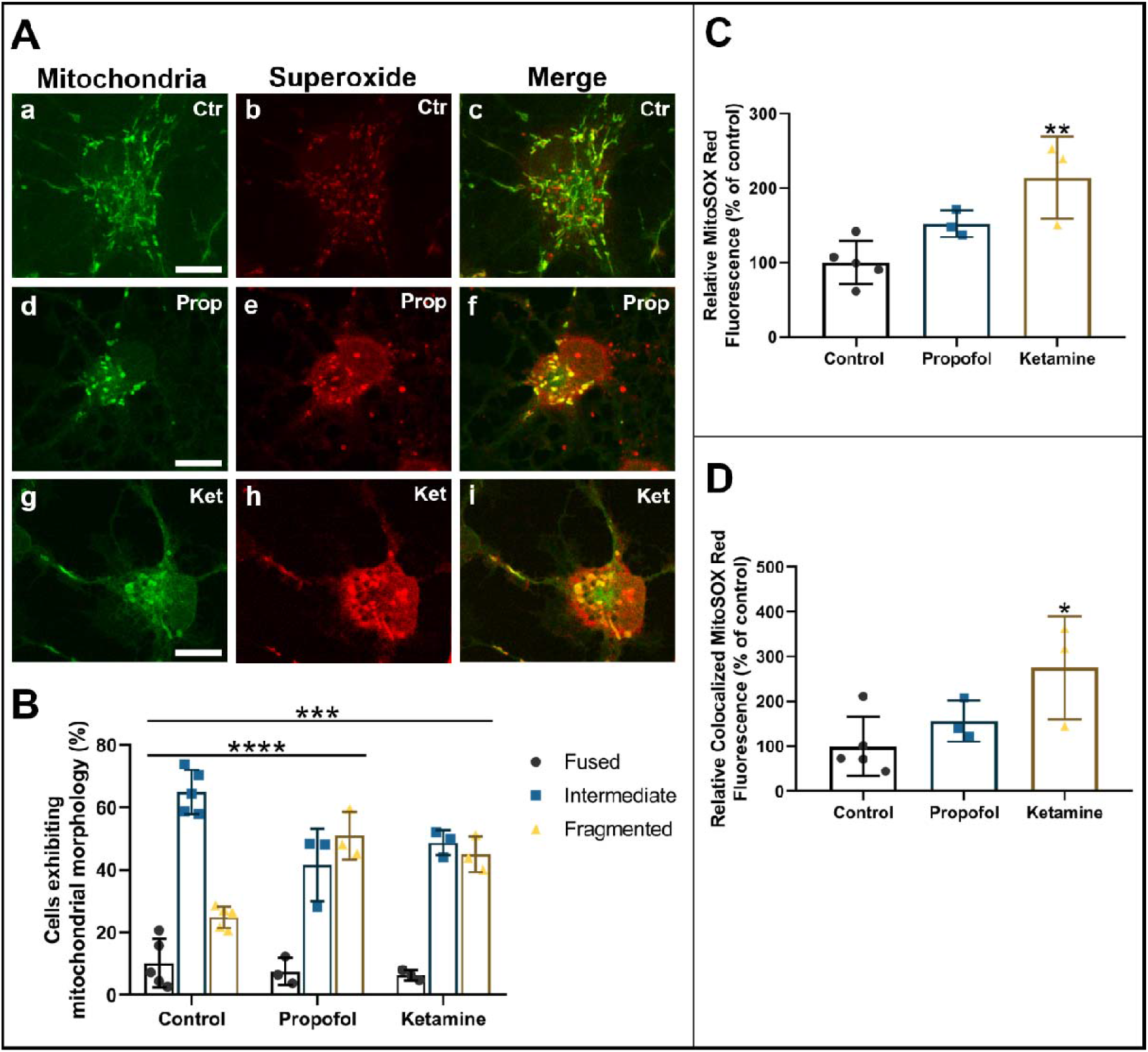
Propofol and ketamine induce mitochondrial fragmentation and ketamine dramatically increases SO production in primary cortical neurons. Representative cropped fluorescent images showing neurons from each condition which were co-stained with MitoTracker™ Green FM for mitochondria and the mitochondrial SO indicator MitoSOX™ Red **(A)**. Quantification of predominant mitochondrial morphology in each treatment. Both propofol and ketamine caused an increase in the proportion of cells exhibiting fragmented mitochondrial networks **(B)**. Quantification of relative SO production **(C)**. Propofol-treated neurons did not significantly differ from control, whereas ketamine more than doubled relative SO production. Quantification of relative SO based on MitoSOX™ Red signal which colocalized to MitoTracker™ Green FM **(D)**. Propofol did not significantly impact mitochondrial SO production, while ketamine almost tripled mitochondrial SO production compared to control. One-way or two-way ANOVA. * *p* < 0.05, ** *p* < 0.01, *** *p* < 0.001, **** *p* < 0.0001 as determined by pairwise comparison of post-hoc tests. Bars indicate ±SD across at least biological replicates. Each data-point indicates the either the percentage of cells that exhibited a specific morphology per dish, or the mean SO signal recorded per dish. Scale bars indicate 10μm.

### A synthetic peptide prevents desflurane-induced mitochondrial fragmentation

Previous studies have attempted to identify protective agents that may mitigate various aspects of anesthetic neurotoxicity. Specifically, we have previously demonstrated that desflurane-induced fragmentation is prevented by pre-treatment of cells with a mitochondrial fission inhibitor, mdivi-1^2^. However, the off-target effects of this drug range from inhibiting cellular proliferation to respiratory impairment and inhibition of basal levels of fission; these negative attributes thus negate its clinical potential and also prevent us from identifying the specific fission protein interactions that may be affected by anesthetics^1,23,24^. Instead, here we used a rationally designed synthetic peptide^25^ that targets excessive “pathological” mitochondrial fission by selectively blocking the dynamin-1-like protein (Drp1) and mitochondrial fission 1 protein (Fis1) interaction, which is upregulated during cellular stress. Blocking only this and not Drp1-receptor interactions that mediate physiological mitochondrial fission was deemed by us to help further elucidate the mechanisms of anesthetic neurotoxicity.

We first investigated whether pre-treatment with the Drp1/Fis1 fission inhibitor P110 could help maintain mitochondrial network integrity both by itself and from desflurane-induced mitochondrial fragmentation. For this test, neurons were either treated with P110 (1μM) mixed in culture media or media alone (control) on the day of cell plating and then exposed either to 4.3 Vol.% desflurane mixed with medical air, or medical air alone for 1 hour. This concentration for P110 corresponds to lower-end dosage reported in literature^25^, and was validated further with tests in our lab **(Figure S1)**. The cells were then maintained in culture and imaged after 4 days using Mitotracker™ Red CMXRos. Control neurons mostly exhibited typical, rod-like morphology between 0.75μm and 3μm in size **(Figure 5a)**. Desflurane-exposed cells, on the other hand, exhibited smaller, more rounded mitochondrial morphologies **(Figure 5b)**. Neurons treated with P110 appeared indistinguishable from control **(Figure 5c and 5d)**. To validate these observations further, we quantified the neurons based on predominant mitochondrial morphologies using a 3-point fragmentation scale (fragmented, intermediate, and fused) **(Figure 1)** and assessed the data via two-way ANOVA. Results were confirmed with post-hoc tests. We found that the cells exposed to clinical levels of desflurane exhibited increased fragmentation (control mean = 22.8%, *SD* = 7.14%, n = 74 cells; desflurane alone mean = 50.67%, *SD* = 1.49%, *p* = 0.005, n = 156 cells). On the other hand, the morphological signatures of cells treated with P110 remained unchanged when compared to control (fragmented: *M* = 18.89%, *SD* = 10.29%, *p* = 0.7674, n = 141 cells). Remarkably, cells pre-treated with P110 and then exposed to desflurane exhibited no significant difference from control or P110 alone (fragmented: *M* = 21.6%, *SD* = 5.6%, *p* = 0.7274 compared to control, *p* = 0.9999 compared to P110 alone, n = 79 cells). The morphological signatures did, however, differ from cells exposed to desflurane alone (*p* = 0.0466) **(Figure 5e)**. These data demonstrate that desflurane increased fragmentation of the mitochondria in primary rat cortical neurons, whereas P110 did not elicit any mitochondrial toxicity by itself. Furthermore, P110 pre-treatment attenuated desflurane-induced fragmentation, thus preserving mitochondrial networks.

**Figure 5:**
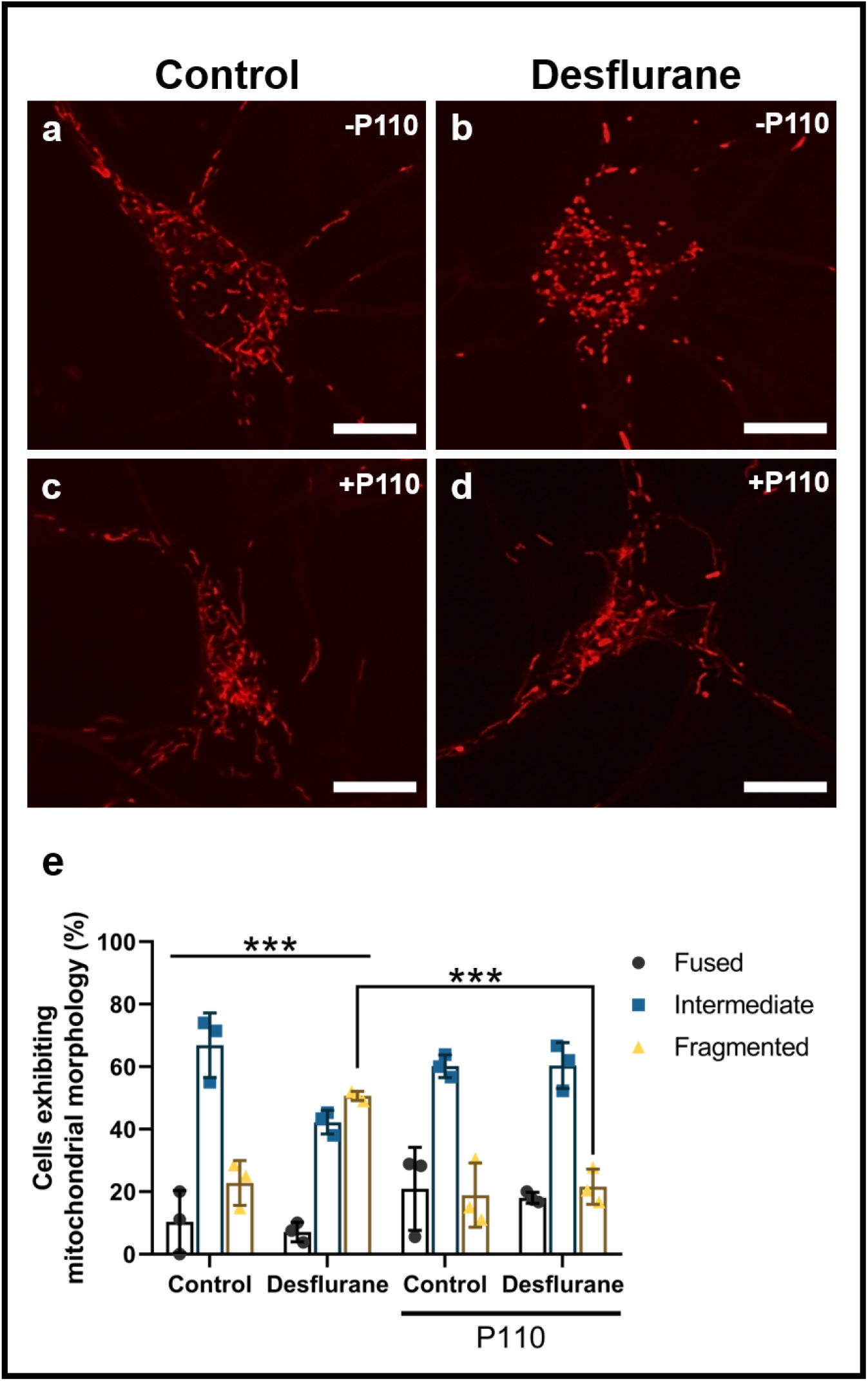
P110 treatment prevents desflurane-induced perturbation of mitochondrial morphology. Fluorescent cropped images showing representative neurons from each condition **(a-d)**. Quantification of predominant mitochondrial morphology in each treatment **(e)**. Desflurane alone caused increased fragmentation. P110 alone did not affect mitochondrial morphologies. P110 pre-treatment, followed by desflurane exposure prevented anesthetic-induced aberrations. Two-Way ANOVA. *** *p*◻<◻0.001 as determined by Tukey’s multiple comparisons post-hoc test. Bars indicate ±SD across biological replicates. Each data-point indicates the percentage of cells that exhibited a specific morphology per biological replicate. Scale bars indicate 10μm.

### The synthetic peptide P110 reverses propofol-induced mitochondrial fragmentation but not ketamine-induced fragmentation and superoxide production

We further examined whether pre-treatment with the Drp1/Fis1 fission inhibitor P110 could help mitigate any changes in SO production and mitochondrial morphology caused by propofol and ketamine. We have shown above that both propofol and ketamine cause excessive mitochondrial fragmentation. We observed that neurons treated with P110 alone did not show any differences from control (fragmented: *M* = 28.04%, *SD* = 3.85%, *p* = 0.9819, n = 144 cells). Remarkably, P110 pre-treatment prevented excessive mitochondrial fragmentation caused by propofol and preserved the overall morphological signature as seen in control (*M* = 31.02%, *SD* = 1.82%, *p* = 0.9170 compared to control, *p* = 0.9923 compared to P110 alone, *p* = 0.01 compared to propofol alone, n = 147 cells) **(Figure 6B)**. Neurons pre-treated with P110 appeared as a mixture between control and fragmented conditions. Specifically, although these neurons appeared less fragmented than those treated with ketamine alone, they were not however, at the level of the controls, nor were they significantly different from ketamine alone (*M* = 34.94%, *SD* = 6.72%, *p* = 0.3464 compared to control, *p* = 0.7697 compared to P110 alone, *p* = 0.4743 compared to ketamine alone, n = 77 cells). Thus, the overall morphological signature still showed a trend towards fragmentation, although not to the level of ketamine alone **(Figure 6B)**. Taken together, these data demonstrate that pre-treatment with the Drp1/Fis1 fission inhibitor P110 protects primary cortical neurons from propofol-induced mitochondrial fragmentation and may attenuate ketamine-induced fragmentation but does not completely reverse morphological changes.

**Figure 6:**
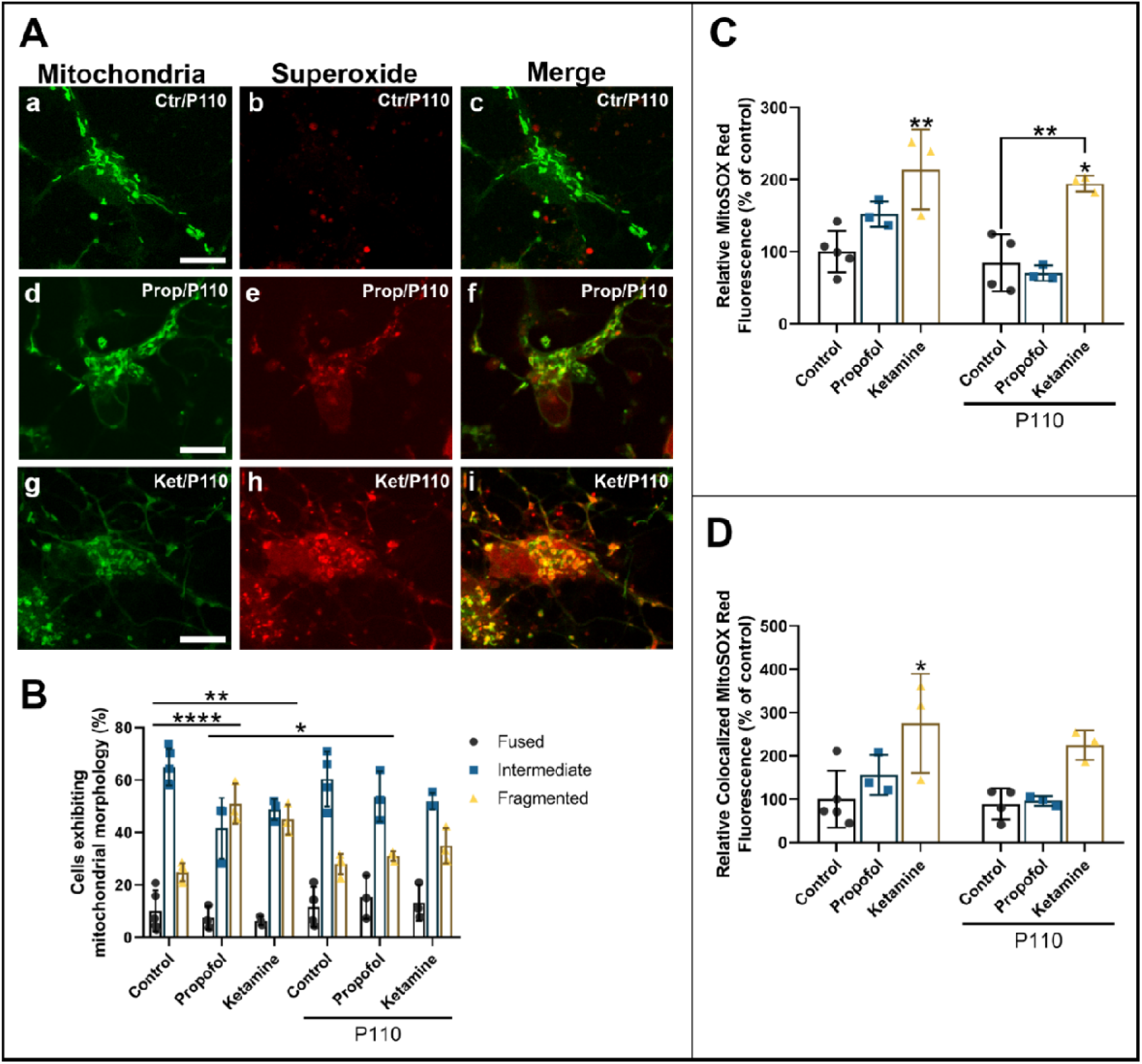
P110 reverses propofol-induced SO production and mitochondrial fragmentation. Representative cropped fluorescent images showing neurons from each condition which are co-stained with MitoTracker™ Green FM for mitochondria and the SO indicator MitoSOX™ Red **(A)**. Quantification of predominant mitochondrial morphology in each treatment **(B)**. For the ease of comparison, the control and anesthetic alone data are recapitulated here. P110 alone did not affect mitochondrial morphology whereas P110 pre-treatment fully rescued against propofol but not ketamine-induced fragmentation. Quantification of relative SO production. For the ease of comparison, the control and anesthetic alone data are recapitulated here **(C)**. P110 alone did not affect SO production, however, P110 pre-treatment completely protected against propofol-induced SO production but not ketamine-induced SO production. Quantification of relative SO based on MitoSOX™ Red signal which colocalized to MitoTracker™ Green FM **(D)**. For the ease of comparison, the control and anesthetic alone data are recapitulated here. P110 reversed propofol-induced elevation in mitochondrial SO production but not ketamine-induced changes. Two-Way ANOVA. * *p* < 0.05, ** *p* < 0.01, **** *p* < 0.0001 as determined by pairwise comparison of post-hoc tests. Bars indicate ±SD across at least biological replicates. Each data-point indicates the either the percentage of cells that exhibited a specific morphology per dish, or the mean SO signal recorded per dish. Scale bars indicate 10μm.

Neurons treated with P110 alone displayed no change in SO levels as compared to control (*M* = 84.30%, *SD* = 39.46% of control, *p* > 0.9999 compared to control, n = 72 areas). Cells treated with P110 followed by propofol exhibited SO expression which was not significantly different from either control, P110 alone, or propofol alone (*M* = 70.20%, *SD* = 10.69% of control, *p* = 0.9762 compared to control, *p* > 0.9999 compared to P110 alone, *p* = 0.0961 compared to propofol alone, n = 68 areas) **(Figure 6C)**. P110 did not reverse ketamine-induced SO production (*M* = 194.48%, *SD* = 10.88% of control, *p* = 0.0155 compared to control, *p* = 0.0061 compared to P110 alone, *p* > 0.9999 compared to ketamine alone, n = 53 areas) **(Figure 6C)**. Of these, the SO signal that colocalized with the mitochondrial signal showed similar trends. P110 by itself did not impact mitochondrial SO (*M* = 88.80%, *SD* = 36.29% of control, *p* = 0.9997, n = 72 areas) compared to control (*M* = 100%, *SD* = 65.42% of control, n = 85 areas). The peptide also did not impact propofol-induced SO production (*M* = 96.34%, *SD* = 11.24% of control, *p* > 0.9999 compared to control, *p* = 0.8202 compared to propofol alone, n = 68 areas). P110 did not significantly impact ketamine-induced SO production, and was not significantly different from control either (*M* = 224.96%, *SD* = 34.43% of control, *p =* 0.1036 compared to control, *p* = 0.0839 compared to P110 alone, *p* = 0.9041 compared to ketamine alone) **(Figure 6D)**. Together, these data demonstrate that the synthetic Drp/Fis1 fission inhibitor P110 did not affect mitochondrial SO production by itself and prevented propofol but not ketamine-induced changes to mitochondrial morphology when administered concomitantly with the tested anesthetic **(Figure 6)**.

### P110 rescues neurons from anesthetic-induced apoptosis

To validate further the neuroprotective properties of P110 against anesthetic-induced apoptosis, we next assessed cell viability of the neuronal cultures. Two-way ANOVA and subsequent post-hoc tests showed that cells treated with P110 alone maintained survival rates similar to those of control (*M* = 91.96%, *SD* = 1.68% alive, *p* = 0.9602 compared to control, n = 42 areas). Remarkably, cells pre-treated with P110 followed by desflurane exposure exhibited no change from control or P110 alone in terms of viable cells and were significantly better than those exposed to desflurane alone (*M* = 92.45%, *SD* = 0.63% alive, *p* = 0.9195 compared to control, *p* > 0.9999 compared to P110 alone, *p* = 0.0018 compared to desflurane alone, n = 51 areas) **(Figure 7; 1d)**. These data demonstrate that exposure to 4.3 Vol.% desflurane results in significant neuronal death *in vitro*, whereas P110 pre-treatment provides neuroprotection against desflurane-induced apoptosis of primary rat cortical neurons.

**Figure 7:**
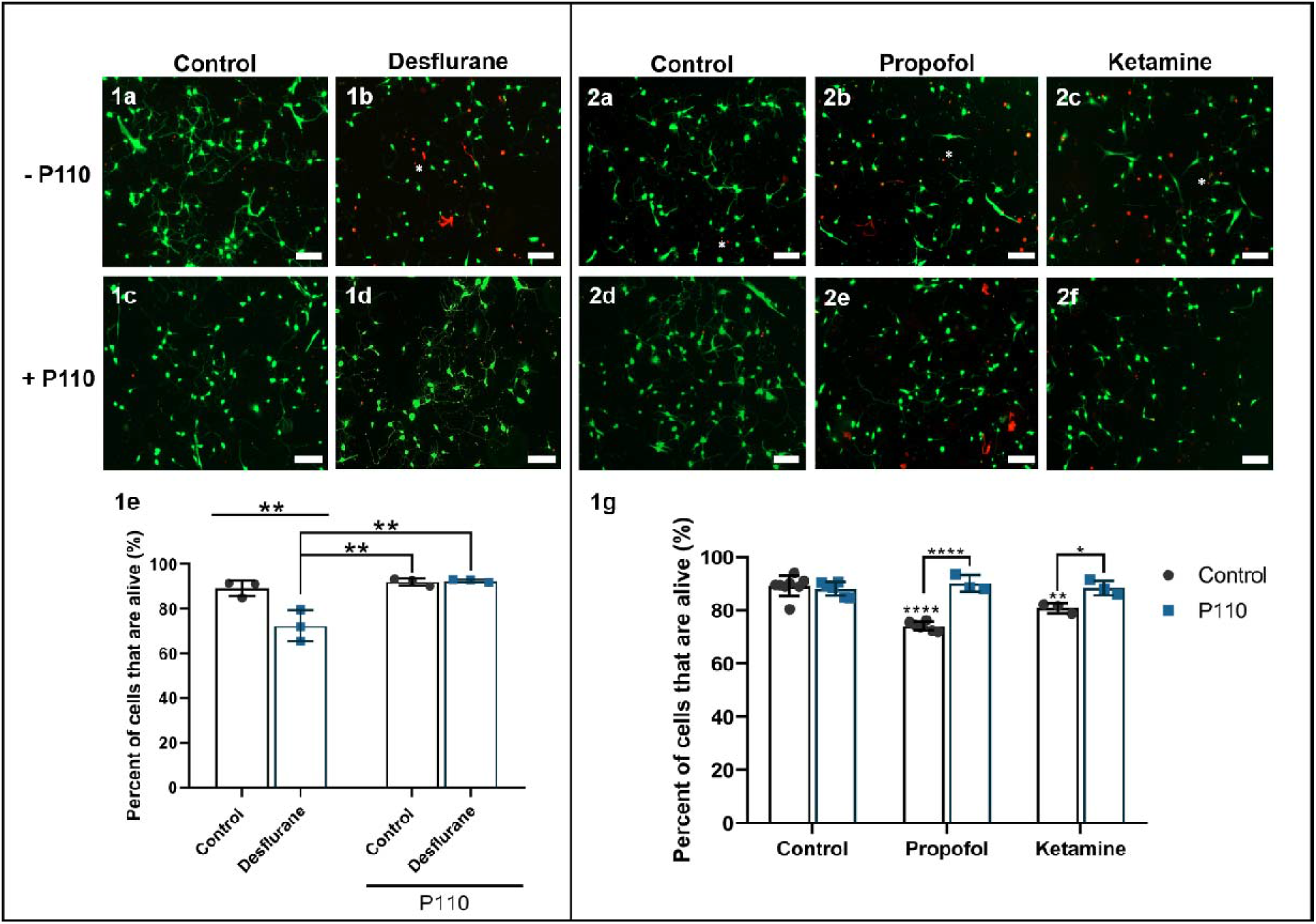
The synthetic peptide P110 rescues neurons from anesthetic-induced apoptosis. Representative live-fluorescent images of different conditions labeled with a LIVE/DEAD™ Viability/Cytotoxicity Assay where live cells are identified with calcein-AM (green) and dead cells are identified with ethidium homodimer-1 (red) **(1a-1d, 2a-2f)**. Quantification of percentage of live cells **(1e,1g)**. For the ease of comparison, the control and anesthetic alone data are recapitulated here. P110 alone did not affect cellular viability whereas P110 pre-treatment protected against desflurane and propofol-induced cell death, whereas P110 pre-treatment followed by ketamine did not differ from control. Two-Way ANOVA. * *p* < 0.05, ** *p* < 0.01, **** *p* < 0.0001 as determined by pairwise comparison of post-hoc tests. Bars indicate ±SD measured across biological replicates. Each data-point indicates the mean viability reported in each biological replicate. Scale bars indicate 100μm.

Similar observations were made for intravenous agents, wherein cells treated with P110 alone maintained cell viability (*M* = 88.13%, *SD* = 2.62%, *p* = 0.9861 compared to control, n = 37 areas). Neurons treated with P110 followed by propofol treatment exhibited no change from those of control or P110 alone, and viability was also significantly higher than cells treated with propofol alone (*M* = 90.15%, *SD* = 3.06% alive, *p* = 0.9953 compared to control, *p* = 0.9198 compared to P110 alone, *p* < 0.0001 compared to propofol alone, n = 15 areas) **(Figure 7; 2e)**. P110 followed by ketamine exposure yielded similar results, reversing ketamine-induced cell death (*M* = 88.45%, *SD* = 2.77% alive, *p* = 0.9992 compared to control, *p* > 0.9999 compared to P110 alone, *p* = 0.0421 compared to ketamine alone, n = 20 areas) **(Figure 7; 2f)**. These data demonstrate that P110 provides protection against the neuro-apoptotic effects of both propofol and ketamine *in vitro.*

### P110 rescues desflurane, propofol, and ketamine-induced effects on neurite outgrowth and synaptogenesis

#### P110 rescues anesthetic-induced changes in cortical neurite outgrowth

To validate the neuroprotective potential of P110, we next tested the effects of desflurane, propofol, and ketamine on the extent of growth after pre-treatment with P110. Cells treated with P110 alone (*M* = 0.96, *SD* = 0.04 fraction of control, *p* = 0.9794 compared to control, n = 73 areas) and those treated with P110 followed by desflurane exposure showed no significant change from control (*M* = 1.01, *SD* = 0.16 fraction of control, *p* = 0.9992 compared to control, *p* = 0.9536 compred to P110 alone, *p* = 0.0553 compared to desflurane alone, n = 54 areas) **(Figure 8A)**. Similarly, we discovered that propofol did not affect neurite outgrowth in developing primary cortical neurons. Neurons treated with P110 alone (*M* = 0.90, *SD* = 0.09 fraction of control, *p* = 0.9908 compared to control, n = 60 areas), or P110 followed by propofol treatment also showed no deviation from control (*M* = 0.95, *SD* = 0.09 fraction of control, *p* > 0.9999 compared to control, *p* > 0.9999 compared to P110 alone, *p* = 0.9962 compared to propofol alone, n = 37 areas) **(Figures 8A)**. These data reveal that propofol does not affect average neurite outgrowth in developing cells.

**Figure 8:**
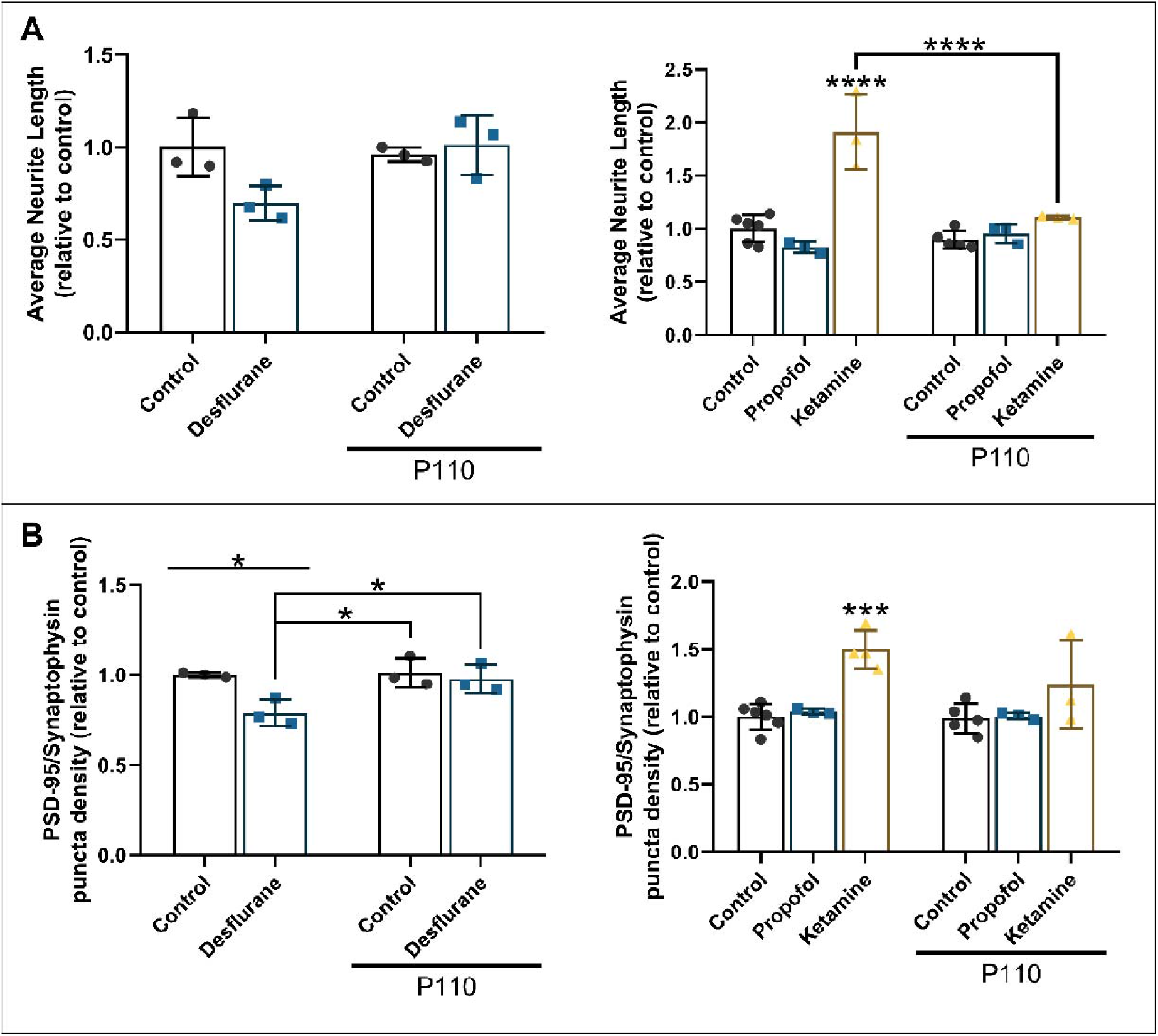
P110 attenuates anesthetic-induced changes in neurite growth and synaptogenesis. Quantification of average neurite length relative to control **(A)**. For the ease of comparison, the control and anesthetic alone data are recapitulated here. P110 alone did not affect neurite outgrowth wheras P110 pre-treatment mitigated ketamine-induced changes. P110 treatment followed by propofol did not differ from control. Quantification of average PSD-95/synaptophysin puncta per neurite length relative to control **(B)**. For the ease of comparison, the control and anesthetic alone data are recapitulated here. P110 alone did not affect synaptic network assembly. P110 pre-treatment mitigated desflurane-induced pertubations and partially prevented ketamine-induced synaptogenesis. P110 followed by propofol treatment did not differ from control. Two-Way ANOVA. * *p* < 0.05, ** *p* < 0.01, *** *p*◻<◻0.001, **** *p* < 0.0001 as determined by pairwise comparison of post-hoc tests. Bars indicate ±SD measured across biological replicates. Each datapoint indicates the mean average neurite lengths, and mean puncta density per biological replicate.

Neurons treated with ketamine exhibited extensive growth compared to control, whereas pre-treatment with P110 attenuated the excessive growth seen in ketamine-treated cells and showed no difference from control (*M* = 1.11, *SD* = 0.02 fraction of control, *p* = 0.9975 compared to control, *p* = 0.6732 compared to P110 alone, *p* < 0.0001 compared to ketamine alone, n = 23 areas) **(Figure 8A)**. Taken together, these data show that ketamine enhances neurite outgrowth of developing cortical neurons, and that these effects are also reversed by P110 pre-treatment.

#### P110 attenuates anesthetic-induced changes in synaptic network assembly in cortical neurons

We next asked whether P110 could provide neuroprotection against anesthetic-induced toxicity regarding its ability to preserve neuronal network health and synaptic assembly. We found that P110 treatment did not affect total synaptic puncta density by itself (*M* = 1.01, *SD* = 0.08 fraction of control, *p*◻= 0.996 compared to control, n = 94 neurites) and instead reversed the desflurane-induced decrease in density of synaptic puncta (*M* = 0.98, *SD* = 0.08 fraction of control, *p*◻= 0.977 compared to control, *p* = 0.9235 compared to P110 alone, *p*◻= 0.0364 compared to desflurane alone, n = 99 neurites) **(Figure 8B)**. Accordingly, P110 treatment alone (*M* = 0.99, *SD* = 0.11 fraction of control, *p*◻> 0.9999 compared to control, n = 97 neurites) or followed by propofol treatment did not change synaptic puncta density compared to control in the intravenous anesthetic tests (*M* = 1.00, *SD* =◻0.03 fraction of control, *p*◻> 0.9999 compared to control, *p* > 0.9999 compared to P110 alone, *p* = 0.9997 compared to propofol alone, n = 68 neurites) **(Figure 8B)**. Interestingly, P110 pre-treatment followed by ketamine exposure did not differ from control cells, or to the synonymous P110 alone cells (*M* = 1.24, *SD* =◻0.33 fraction of control, *p* = 0.2237 compared to control, *p* = 0.2114 compared to P110 alone, *p*◻= 0.2244 compared to ketamine alone, n = 57 neurites) **(Figures 8B)**. Taken together, these data show for the first time, that treatment with the mitochondrial fission inhibitor P110 rescues anesthetic-mediated reduction in synaptic puncta density in cortical neurons, completely reversing desflurane-induced decreases in the number of boutons, and partially preventing ketamine-induced increases.

## Discussion

There is now significant awareness of the potential neurotoxic effects of anesthetic agents on brain development, and progress has been made in understanding the mechanisms underlying the evidence of harm^4,10,20^. However, despite several studies investigating these effects, developing a consensus vis-à-vis the effects of anesthetics on neuronal health has remained elusive. The reasons for this lack of consensus perhaps owe their existence to the fact that myriad models have been subjected to different anesthetic agents and under diverse experimental conditions, thus generating irreconcilable conclusions. As such, neither the precise mechanisms, nor the extent to which these effects may impact any particular patient can be considered reliably. Hence, we deemed it important to resolve these differing conclusions by creating identical experimental conditions through which to test the effects of key inhalational and intravenous general anesthetics on developing neural circuits. We selected this experimental paradigm for two main reasons. The primary culture setup offers an opportunity for a high-quality assessment of fundamental characteristics, from the forming of synaptic networks, to live observation of subcellular characteristics at a period that is critical for neurodevelopment that would have otherwise not been possible. Such an analysis at the cellular and molecular level is not feasible in more intact, freely behaving animals.

Interestingly, we discovered that under identical cell culture conditions employing only cortical neurons, the anesthetics exerted differing effects on neurite growth and synaptic network assembly. While the GABAergic anesthetic agent desflurane decreased synaptic density, the NMDAergic ketamine had the opposite effect, whereas propofol had no significant impact on these parameters. This suggests an agent-specific influence on key characteristics of neuronal development. Even with these differing effects however, all three anesthetics caused significant cell death, underscoring the neuro-apoptotic nature of these agents. Although we showed morphological, and not functional attributes of a synapse, our previous experiments – conducted under identical conditions – have nevertheless demonstrated that decreased synaptic puncta are always corroborated with lower amplitude and frequency of spontaneous synaptic events^2^. The differences observed between agents thus support and build on previous studies that have investigated these drugs individually and substantiate the notion that regardless of the model system used, both inhalational and intravenous general anesthetics exert different effects on neuronal viability and growth.

For example, previous studies have demonstrated propofol-induced acute neurite retraction in primary rat cortical neurons (minutes after anesthetic exposure) which was completely reversed within 10 minutes after the washout for lower concentrations (0.02, 0.2, and 2μM)^26^. Although these live-imaging tests are consistent with our observations, the authors noted that the effects of longer exposure times or exposure to 20 μM propofol could not be reversed. This may be explained by the short timeline of the experiment, with neurite lengths being observed within hours of anesthetic exposure, and not days afterwards to quantify lasting effects. For the impact of propofol on synaptogenesis, divergent results have been reported with regards to cell type and the age of animals^27,28^. Similarly, ketamine has been previously shown to either downregulate synaptic proteins in the hippocampus of the developing brain^29^, or to upregulate them in the cortex^30^. A 2010 study demonstrated that activation of the mammalian target of rapamycin pathway underlies ketamine-induced synaptogenesis by illustrating increased dendritic spine density in cortical neurons. This was coupled with a functional assessment of synaptic activity in pyramidal cells^31^.

However, consolidating differences between previous studies cannot explain the variations observed between these anesthetics vis-à-vis their relative impacts of key aspects of neuronal development. To help elucidate this, and to identify a potential common mechanism underlying anesthetic-induced neurotoxicity in developing neurons, we directly investigated neuronal energy dynamics, specifically mitochondrial structure and function. In recent years, the roles of disrupted mitochondrial networks and compromised mitochondrial dynamics have also come to light in several neurodegenerative diseases, although clinical treatments remain limited^32^. Changes in cellular growth machinery and cytoskeletal protein function heavily involve these networks; and the delivery of functional mitochondria to sites with high metabolic requirements (such as growth cones and branch points) is crucial in synapse formation and synaptic transmission. Thus, any perturbation in itochondrial dynamics may have lasting implications on neuronal network development^33^. We speculate that this may explain the trends observed in our results and the differences in the field with regards to the comparison between the primarily NMDAergic ketamine and the GABAergic propofol.

The widely documented GABA depolarizing-to-hyperpolarizing polarity switch in the developing brain^34,35^ may underlie the differences in the effects of these agents. GABAergic transmission plays an important role in propagating giant depolarizing potentials, which are crucial for early brain development and for additional GABAergic synaptogenesis^34,36^. We speculate that anesthetic-induced GABA agonism by agents such as isoflurane^37^, desflurane^38^, propofol^39,40^, and to a minor degree, ketamine^41^, may be a factor responsible for increasing GABAergic sensitivity in developing brains when administered to pregnant mother or neonates. In the context of apoptosis, this may underlie anesthetic-induced GABAergic excitotoxicity^37^ and thus, cell death. Acting in opposition to this, ketamine-mediated NMDA antagonism^42^ may counteract these effects, and provide protection against excitotoxicity^43^. Thus, various reports suggesting ketamine’s neuroprotective role in brains exposed to noxious stimuli cannot be discounted when investigating anesthetic neurotoxicity^44^. For instance, a 2018 study found that ketamine prevented behavioral deficits caused by isoflurane^45^, an agent that induces GABAergic excitotoxicity in the developing brain^37^. Preventing hyper-active synaptic transmission which may initiate cell death and modulating mitochondrial function may complement ketamine’s mTOR-mediated anti-depressant properties^31^ in what would have otherwise been a highly excitotoxic cellular environment.

This dual function may also explain ketamine’s purported neuroprotective effects in the developing brain. Various investigators have commented on the seemingly opposite effects of ketamine and have correctly suggested that drug dose, frequency of exposure, and the noxious state of the brain may impact ketamine-induced neurotoxicity and neuroprotection^46^. We believe this mechanistic understanding may provide a clearer answer to these exciting set of questions, many of which remain elusive. For example, determining whether ketamine could by itself relieve anesthetic-induced excitotoxicity may help explain its clinical uses. These insights could offer a unique opportunity to develop novel strategies to mitigate anesthetic-induced neurotoxicity in the clinic.

In search for such strategies, we previously demonstrated that both desflurane and sevoflurane-induced neurotoxicity could be reversed by the fission inhibitor mdivi-1 using the same experimental paradigm^2^. A 2015 study using a cell line showcased similar results for propofol, which too, were attenuated by mdivi-1^47^. However, the clinical applications of mdivi-1 are limited due to its off-site effects^48^. Instead, selectively blocking Drp1/Fis1 interaction while not affecting other physiological interactions could specifically inhibit excessive fission in stressed conditions such as anesthetic exposure. We showed here that using P110, a non-toxic, seven amino acid long selective inhibitor of Drp1/Fis1 provides protection and complete rescue from both desflurane and propofol-induced neurotoxicity in primary cortical neurons, and partial recovery from ketamine-induced changes. These results are the first to demonstrate the protective effects of P110 against anesthetic-induced neurotoxicity. To our knowledge this is also the first study to link Fis1 receptor-mediated fission with anesthetic-induced aberrations. Since the peptide protects against both GABAergic and NMDAergic agents (although there is some overlap in action^4^), this suggests a common, activity-based protection via the mitochondria which may have resulted from a myriad of factors; including changes in cellular energetics, hyperactivity, or other still-to-be explored interactions of anesthetics with mitochondria. Although we have brought together a few crucial pieces towards characterizing mechanistic interactions between anesthetics and mitochondrial health, further work to identify how key mitochondrial cycles are regulated by anesthetics requires attention. Investigating whether mitochondrial fission inhibitors such as P110 can mitigate anesthetic-induced effects *in vivo* also remains to be determined and will be the focus of future work.

Taken together, anesthetic-induced neurotoxicity is a significant issue. However, given that anesthetics are essential tools that enable critical surgeries, they are unavoidable. This does not however, mean that their cytotoxic and long-term effects should be overlooked as it would undermine our efforts towards searching for least toxic anesthetic agents or to synthesize compounds that would mitigate their cytotoxic effects in humans.

This study makes two important contributions to our understanding of anesthetics and neurotoxicity. First, we provide a pragmatic account of both commonly used inhalational and intravenous general anesthetic agents with different modes of action in relation to their effects on key aspects of neuronal health under identical experimental conditions. We propose a mechanistic-based paradigm to the varying results observed in the field. We integrate these findings with current interpretations of anesthetic-induced neurotoxicity and attempt to fill several gaps in our understanding of anesthetic-induced harm in cultured rat cortical neurons. Second, we identified a potential, novel protective strategy against anesthetic-induced neurotoxicity by preserving mitochondrial health. This approach could help design future neuroprotective drugs with potential clinical applications. We show that the neurotoxic effects of the anesthetics are reversed by a selective mitochondrial fission inhibitor, P110, showcasing tremendous *in vivo* and clinical potential. The findings presented here thus underscore the fact that preserving mitochondrial networks may be key for future research into potential neuroprotective agents that will complement the vital role of anesthetics in the operating theatre.

## Additional Information

### Author Contributions

F.I., J.S., and N.I. designed the experiments. F.I., A.T., U.A., M.P, K.J., N.JT. R.S. and S.B performed the experiments. F.I., A.T., U.A., M.P, K.J., A.I., and J.C. anonymized, collected, and analyzed the data, P.M. contributed the mitochondrial SO expression protocol. F.I. and N.I. wrote and edited the manuscript with editorial input from A.T., U.A., M.P, K.J., N.JT. R.S., J.C., P.M., T.S., K.Y., J.S., and T.R. All authors approved the final version of the manuscript and agree to be accountable for all aspects of the work in ensuring that questions related to the accuracy or integrity of any part of the work are appropriately investigated and resolved. All persons designated as authors qualify for authorship, and all those who qualify for authorship are listed.

## Acknowledgements

We would like to thank Jean Kawasoe and Wali Zaidi for their excellent technical support and Daria Mochly-Rosen for providing the P110 peptide for the experiments.

## Supplementary Material

**Figure S1:**
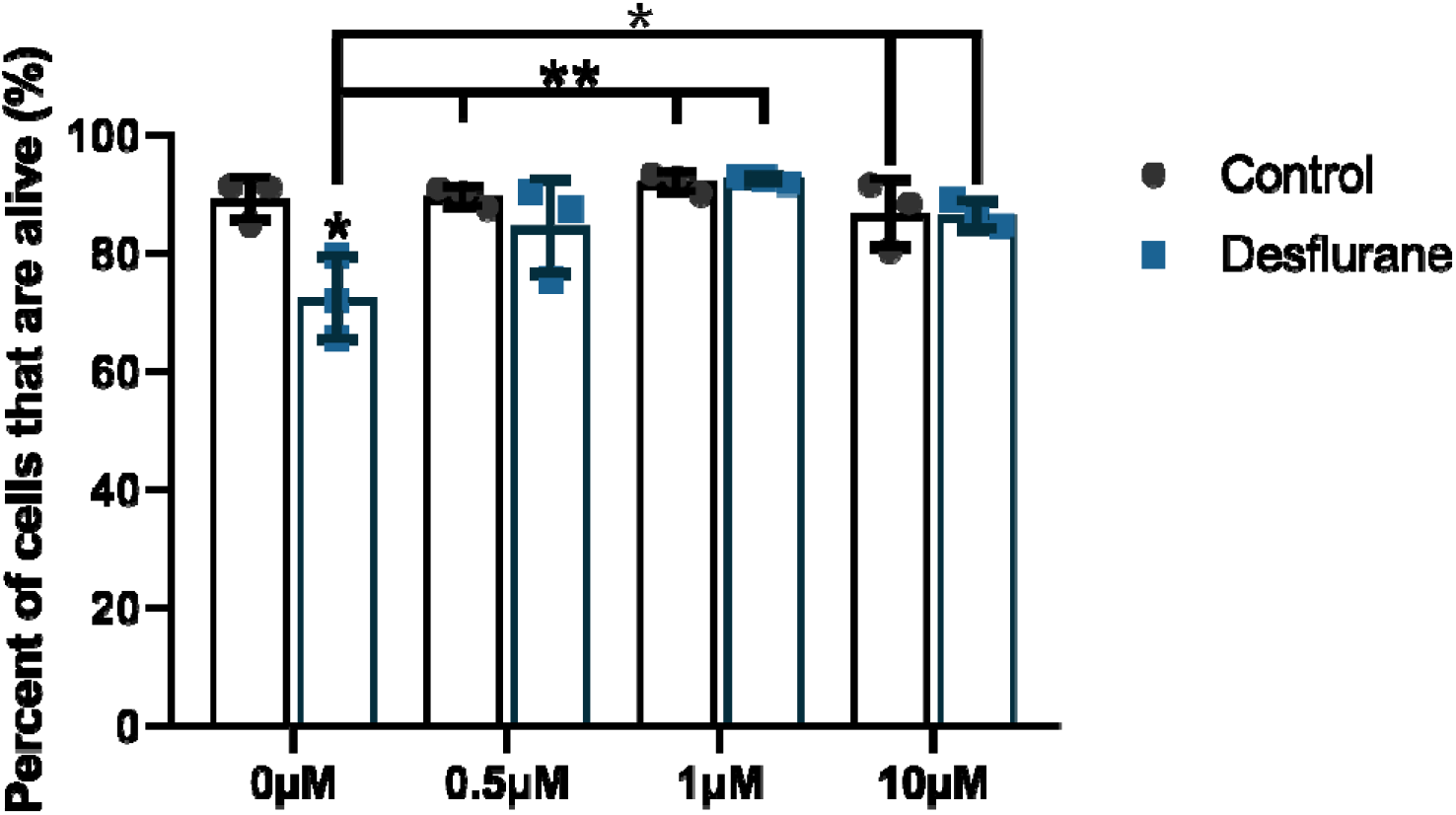
P110 prevents desflurane-induced cell death in a concentration-dependent manner. P110 did not affect cell viability by itself (0.5μM: *M* = 89.48%, *SD* = 1.67% alive, *p* > 0.9999 compared to control, n = 37 areas; 1μM: *M* = 91.96%, *SD* = 1.68% alive, *p* > 0.9999 compared to control, n = 42 areas; 10μM: *M* = 86.74%, *SD* = 5.69% alive, *p* > 0.9999 compared to control, n = 40 areas) and instead reversed desflurane-induced cell death (0.5μM: *M* = 84.43%, *SD* = 7.9% alive, *p* = 0.9993 compared to control, *p* = 0.0089 compared to desflurane alone, n = 36 areas; 1μM: *M* = 92.45%, *SD* = 0.63% alive, *p* > 0.9999 compared to control, *p* = 0.0018 compared to desflurane alone, n = 45 areas; 10μM: *M* = 86.4%, *SD* = 2.34% alive, *p* > 0.9999 compared to control, *p* = 0.0402 compared to desflurane alone, n = 66 areas). Two-Way ANOVA. * *p* < 0.05, ** *p* < 0.01. Experiments and results were validated at least twice. Bars indicate ±SD across biological replicates. Each data-point indicates the mean viability reported in each biological replicate.

## References

1 Dokkedal, U., Hansen, T. G., Rasmussen, L. S., Mengel-From, J. & Christensen, K. Cognitive Functioning after Surgery in Middle-aged and Elderly Danish Twins. Anesthesiology 124, 312–321, doi:10.1097/ALN.0000000000000957 (2016).

2 Xu, F. et al. The mitochondrial division inhibitor Mdivi-1 rescues mammalian neurons from anesthetic-induced cytotoxicity. Mol Brain 9, 35, doi:10.1186/s13041-016-0210-x (2016).

3 Bai, X., Twaroski, D. & Bosnjak, Z. J. Modeling anesthetic developmental neurotoxicity using human stem cells. Semin Cardiothorac Vasc Anesth 17, 276–287, doi:10.1177/1089253213495923 (2013).

4 Iqbal, F. et al. Anesthetics: From modes of action to unconsciousness and neurotoxicity. J Neurophysiol, doi:10.1152/jn.00210.2019 (2019).

5 Flick, R. P. et al. Cognitive and behavioral outcomes after early exposure to anesthesia and surgery. Pediatrics 128, e1053–1061, doi:10.1542/peds.2011-0351 (2011).

6 FDA. FDA drug safety communication: FDA approves label changes for use of general anesthetic and sedation drugs in young children, <https://www.fda.gov/Drugs/DrugSafety/ucm554634.htm> (2017).

7 Canada, H. New warnings regarding the use of certain sedative and anesthetic drugs during pregnancy and in early childhood, <https://healthycanadians.gc.ca/recall-alert-rappel-avis/hc-sc/2017/65516a-eng.php> (2017).

8 Jevtovic-Todorovic, V. et al. Anaesthetic neurotoxicity and neuroplasticity: an expert group report and statement based on the BJA Salzburg Seminar. Br J Anaesth 111, 143–151, doi:10.1093/bja/aet177 (2013).

9 Sun, L. Early childhood general anaesthesia exposure and neurocognitive development. Br J Anaesth 105 Suppl 1, i61–68, doi:10.1093/bja/aeq302 (2010).

10 Jevtovic-Todorovic, V. General Anesthetics and Neurotoxicity: How Much Do We Know? Anesthesiology clinics 34, 439–451, doi:10.1016/j.anclin.2016.04.001 (2016).

11 Joshi, A. U., Saw, N. L., Shamloo, M. & Mochly-Rosen, D. Drp1/Fis1 interaction mediates mitochondrial dysfunction, bioenergetic failure and cognitive decline in Alzheimer’s disease. Oncotarget 9, 6128–6143, doi:10.18632/oncotarget.23640 (2018).

12 Manczak, M., Calkins, M. J. & Reddy, P. H. Impaired mitochondrial dynamics and abnormal interaction of amyloid beta with mitochondrial protein Drp1 in neurons from patients with Alzheimer’s disease: implications for neuronal damage. Hum Mol Genet 20, 2495–2509, doi:10.1093/hmg/ddr139 (2011).

13 Guo, X. et al. Inhibition of mitochondrial fragmentation diminishes Huntington’s disease-associated neurodegeneration. J Clin Invest 123, 5371–5388, doi:10.1172/jci70911 (2013).

14 Park, J.-S., Davis, R. L. & Sue, C. M. Mitochondrial Dysfunction in Parkinson’s Disease: New Mechanistic Insights and Therapeutic Perspectives. Current neurology and neuroscience reports 18, 21–21, doi:10.1007/s11910-018-0829-3 (2018).

15 Sumi, C. et al. Propofol induces a metabolic switch to glycolysis and cell death in a mitochondrial electron transport chain-dependent manner. PloS one 13, e0192796–e0192796, doi:10.1371/journal.pone.0192796 (2018).

16 Venancio, C. et al. Acute ketamine impairs mitochondrial function and promotes superoxide dismutase activity in the rat brain. Anesth Analg 120, 320–328, doi:10.1213/ane.0000000000000539 (2015).

17 Boscolo, A. et al. Early exposure to general anesthesia disturbs mitochondrial fission and fusion in the developing rat brain. Anesthesiology 118, 1086–1097, doi:10.1097/ALN.0b013e318289bc9b (2013).

18 Grundy, D. Principles and standards for reporting animal experiments in The Journal of Physiology and Experimental Physiology. J Physiol 593, 2547–2549, doi:10.1113/jp270818 (2015).

19 AVMA. Vol. 9 62 (National Institute of Health, Office of Intramural Research, Office of Animal Care and Use, National Institute of Health Office of Animal Care and Use, 2020).

20 Berndt, N. et al. Possible neurotoxicity of the anesthetic propofol: evidence for the inhibition of complex II of the respiratory chain in area CA3 of rat hippocampal slices. Arch Toxicol 92, 3191–3205, doi:10.1007/s00204-018-2295-8 (2018).

21 Bosnjak, Z. J. et al. Ketamine induces toxicity in human neurons differentiated from embryonic stem cells via mitochondrial apoptosis pathway. Curr Drug Saf 7, 106–119 (2012).

22 Xu, F., Piett, C., Farkas, S., Qazzaz, M. & Syed, N. I. Silver nanoparticles (AgNPs) cause degeneration of cytoskeleton and disrupt synaptic machinery of cultured cortical neurons. Mol Brain 6, 29, doi:10.1186/1756-6606-6-29 (2013).

23 Wang, X. et al. Alpha-lipoic acid prevents bupivacaine-induced neuron injury in vitro through a PI3K/Akt-dependent mechanism. Neurotoxicology 31, 101–112, doi:10.1016/j.neuro.2009.10.010 (2010).

24 Rosdah, A. A., J, K. H., Delbridge, L. M., Dusting, G. J. & Lim, S. Y. Mitochondrial fission - a drug target for cytoprotection or cytodestruction? Pharmacol Res Perspect 4, e00235, doi:10.1002/prp2.235 (2016).

25 Qi, X., Qvit, N., Su, Y. C. & Mochly-Rosen, D. A novel Drp1 inhibitor diminishes aberrant mitochondrial fission and neurotoxicity. Journal of cell science 126, 789–802, doi:10.1242/jcs.114439 (2013).

26 Turina, D., Loitto, V. M., Bjornstrom, K., Sundqvist, T. & Eintrei, C. Propofol causes neurite retraction in neurones. Br J Anaesth 101, 374–379, doi:10.1093/bja/aen185 (2008).

27 Milanovic, D. et al. Neonatal Propofol Anesthesia Changes Expression of Synaptic Plasticity Proteins and Increases Stereotypic and Anxyolitic Behavior in Adult Rats. Neurotox Res 32, 247–263, doi:10.1007/s12640-017-9730-0 (2017).

28 De Roo, M. et al. Anesthetics rapidly promote synaptogenesis during a critical period of brain development. PloS one 4, e7043, doi:10.1371/journal.pone.0007043 (2009).

29 Li, Y. et al. Long-term neurocognitive dysfunction in offspring via NGF/ ERK/CREB signaling pathway caused by ketamine exposure during the second trimester of pregnancy in rats. Oncotarget 8, 30956–30970, doi:10.18632/oncotarget.16042 (2017).

30 Zhao, T. et al. Prenatal ketamine exposure causes abnormal development of prefrontal cortex in rat. Sci Rep 6, 26865, doi:10.1038/srep26865 (2016).

31 Li, N. et al. mTOR-dependent synapse formation underlies the rapid antidepressant effects of NMDA antagonists. Science (New York, N.Y.) 329, 959–964, doi:10.1126/science.1190287 (2010).

32 Burte, F., Carelli, V., Chinnery, P. F. & Yu-Wai-Man, P. Disturbed mitochondrial dynamics and neurodegenerative disorders. Nature reviews. Neurology 11, 11–24, doi:10.1038/nrneurol.2014.228 (2015).

33 Flippo, K. H. & Strack, S. Mitochondrial dynamics in neuronal injury, development and plasticity. Journal of cell science 130, 671–681, doi:10.1242/jcs.171017 (2017).

34 Ben-Ari, Y. Excitatory actions of gaba during development: the nature of the nurture. Nature Reviews Neuroscience 3, 728–739, doi:10.1038/nrn920 (2002).

35 Represa, A. & Ben-Ari, Y. Trophic actions of GABA on neuronal development. Trends Neurosci 28, 278–283, doi:10.1016/j.tins.2005.03.010 (2005).

36 Spitzer, N. C. How GABA generates depolarization. J Physiol 588, 757–758, doi:10.1113/jphysiol.2009.183574 (2010).

37 Zhao, Y. L. et al. GABAergic excitotoxicity injury of the immature hippocampal pyramidal neurons’ exposure to isoflurane. Anesthesia and analgesia 113, 1152–1160, doi:10.1213/ANE.0b013e318230b3fd (2011).

38 Nishikawa, K. & Harrison, N. L. The actions of sevoflurane and desflurane on the gamma-aminobutyric acid receptor type A: effects of TM2 mutations in the alpha and beta subunits. Anesthesiology 99, 678–684 (2003).

39 Krasowski, M. D. et al. Propofol and other intravenous anesthetics have sites of action on the gamma-aminobutyric acid type A receptor distinct from that for isoflurane. Mol Pharmacol 53, 530–538 (1998).

40 Yip, G. M. S. et al. A propofol binding site on mammalian GABAA receptors identified by photolabeling. Nature Chemical Biology 9, 715–720, doi:10.1038/nchembio.1340 (2013).

41 Wang, D. S., Penna, A. & Orser, B. A. Ketamine Increases the Function of gamma-Aminobutyric Acid Type A Receptors in Hippocampal and Cortical Neurons. Anesthesiology 126, 666–677, doi:10.1097/aln.0000000000001483 (2017).

42 Zorumski, C. F., Izumi, Y. & Mennerick, S. Ketamine: NMDA Receptors and Beyond. J Neurosci 36, 11158–11164, doi:10.1523/jneurosci.1547-16.2016 (2016).

43 Gonzales, Jerry M., MD, Loeb, Alex L., PhD, Reichard, Peter S., MD & Irvine, S., BS. Ketamine Inhibits Glutamate-, N-Methyl-D-Aspartate-, and Quisqualate-stimulated cGMP Production in Cultured Cerebral Neurons Anesthesiology: The Journal of the American Society of Anesthesiologists 82, 205–213 (1995).

44 Anand, K. J. et al. Ketamine reduces the cell death following inflammatory pain in newborn rat brain. Pediatr Res 62, 283–290, doi:10.1203/PDR.0b013e3180986d2f (2007).

45 Wang, R., Zhang, Z., Kumar, M., Xu, G. & Zhang, M. Neuroprotective potential of ketamine prevents developing brain structure impairment and alteration of neurocognitive function induced via isoflurane through the PI3K/AKT/GSK-3beta pathway. Drug design, development and therapy 13, 501–512, doi:10.2147/dddt.s188636 (2019).

46 Yan, J. & Jiang, H. Dual effects of ketamine: neurotoxicity versus neuroprotection in anesthesia for the developing brain. J Neurosurg Anesthesiol 26, 155–160, doi:10.1097/ana.0000000000000027 (2014).

47 Twaroski, D. M. et al. Altered Mitochondrial Dynamics Contributes to Propofol-induced Cell Death in Human Stem Cell-derived Neurons. Anesthesiology 123, 1067–1083, doi:10.1097/aln.0000000000000857 (2015).

48 Bordt, E. A. et al. The Putative Drp1 Inhibitor mdivi-1 Is a Reversible Mitochondrial Complex I Inhibitor that Modulates Reactive Oxygen Species. Dev Cell 40, 583–594.e586, doi:10.1016/j.devcel.2017.02.020 (2017).

